# Inhibiting ribosome assembly and ribosome translation have distinctly different effects on the abundance and paralogue composition of ribosomal protein mRNAs in *Saccharomyces cerevisiae*

**DOI:** 10.1101/2022.11.09.515899

**Authors:** Md Shamsuzzaman, Nusrat Rahman, Brian Gregory, Ananth Bommakanti, Janice M Zengel, Vincent M Bruno, Lasse Lindahl

**Affiliations:** Department of Biological Sciences, University of Maryland Baltimore County, 1000 Hilltop Circle, Baltimore, Maryland 21250, USA; Institute for Genome Sciences, Health Sciences Facility III, 670 West Baltimore St, Baltimore 21201, Maryland, USA

**Author notes:** Md Shamsuzzaman, Bristol Myers Squibb, Lawrenceville, NJ 08648, USA. Nusrat Rahman, Current address: American Psychiatric Association, 800 Maine Avenue, SW, Washington, DC 20024, USA. Ananth Bommakanti. Twist Biosciences Singapore, 3 Fusionopolis Place #03-50 Galaxis (work Loft) Singapore 138523.

## Abstract

Many mutations in genes for ribosomal proteins and assembly factors cause cell stress and altered cell fate resulting in congenital diseases, collectively called ribosomopathies. Even though all such mutations depress the cell’s protein synthesis capacity, they generate many different phenotypes, suggesting that the diseases are not due simply to insufficient protein synthesis capacity. To learn more, we have investigated how the global transcriptome in *Saccharomyces cerevisiae* responds to reduced protein synthesis generated in two different ways: abolishing the assembly of new ribosomes or inhibiting ribosomal function. Our results show that the mechanism by which protein synthesis is obstructed affects the ribosomal protein transcriptome differentially: ribosomal protein mRNA abundance increases during the abolition of ribosome formation but decreases during the inhibition of ribosome function. Interestingly, the ratio between mRNAs from some, but not all, paralogous genes encoding slightly different versions of a given r-protein change differently during the two types of stress, suggesting that specific ribosomal protein paralogues may contribute to the stress response. Unexpectedly, the abundance of transcripts for ribosome assembly factors and translation factors remains relatively unaffected by the stresses. On the other hand, the state of the translation apparatus does affect cell physiology: mRNA levels for some other proteins not directly related to the translation apparatus also change differentially, though not coordinately with the r-protein genes, in response to the stresses.

**Importance:** Mutations in genes for ribosomal proteins or assembly factors cause a variety of diseases called ribosomopathies. These diseases are typically ascribed to a reduction in the cell’s capacity for protein synthesis. Paradoxically, ribosomal mutations result in a wide variety of disease phenotypes, even though they all reduce protein synthesis. Here we show that the transcriptome changes differently depending on how the protein synthesis capacity is reduced. Most strikingly, inhibiting ribosome formation and ribosome function have opposite effects on the abundance of mRNA for ribosomal proteins, while genes for ribosome translation and assembly factors show no systematic responses. Thus, the process by which the protein synthesis capacity is reduced contributes decisively to global mRNA composition. This emphasis on process is a new concept in understanding ribosomopathies and other stress responses.

## Introduction

Eukaryotic ribosome formation involves synthesizing and assembling four rRNA molecules with 79-80 unique ribosomal proteins (r-proteins). More than 250 proteins and ncRNA molecules are required for this complex process (1-4). The number of ribosomes per cell (∼10^5^ in yeast, ∼10^7^ in humans) and their structural complexity make ribosome formation very resource intensive. Mechanisms have evolved to match ribosome biogenesis to growth conditions and growth rate (5-7). For example, the expression of rRNA and r-protein genes is repressed during nutritional insufficiency while genes encoding proteins needed to adapt to the stress condition are upregulated (8, 9). The nutrient shortage inactivates TORC1, a major gatekeeper of ribosome gene transcription, resulting in the replacement of positive transcription factors on ribosomal genes with negative ones. The key role of ribosome metabolism in general cell physiology is also illustrated by the signals produced during the distortion of normal ribosome function which may result in apoptosis and cell death (10-13).

Mutations in numerous genes for r-proteins or assembly factors cause a variety of congenital human diseases, collectively called “ribosomopathies” (14-17). It is assumed that the root for these calamities is the decreased translation capacity, which in turn can lead to the formation of “specialized ribosomes” with altered preferences for specific mRNAs, the interaction of free r-proteins with p53 and cell cycle regulators, and changes to specific metabolic pathways (18-23). One of the standing puzzles of ribosomopathies is that, although mutations in different ribosomal genes all decrease the rate of ribosome formation they generate a variety of distinct disease phenotypes, and, vice versa, mutations in different r-protein genes can cause the same disease, e.g., Diamond Blackfan Anemia (24).

To shed light on the differential effects of assaulting the cell’s capacity for protein synthesis, we used *Saccharomyces cerevisiae* (yeast) to compare the global transcriptomes after reducing the protein synthesis capacity by two different assaults: (i) blocking ribosome formation by abrogating the synthesis of r-protein ribosome uL4 (“nucleolar stress”, also called “ribosome stress”), and (ii) abolishing ribosome function by stopping the synthesis of Translation Elongation Factor eEF3 (“translation stress”). Both acts lower the translation capacity but in completely different ways. Blocking uL4 synthesis reduces the accumulation of newly synthesized ribosomes without affecting the ability of pre-existing ribosomes to translate mRNA. Conversely, abrogating eEF3 synthesis does not affect ribosome number, but progressively inhibits ribosome function (Fig.1). Our results show that the two stresses have profoundly different effects on the transcriptome.

**Fig. 1.**
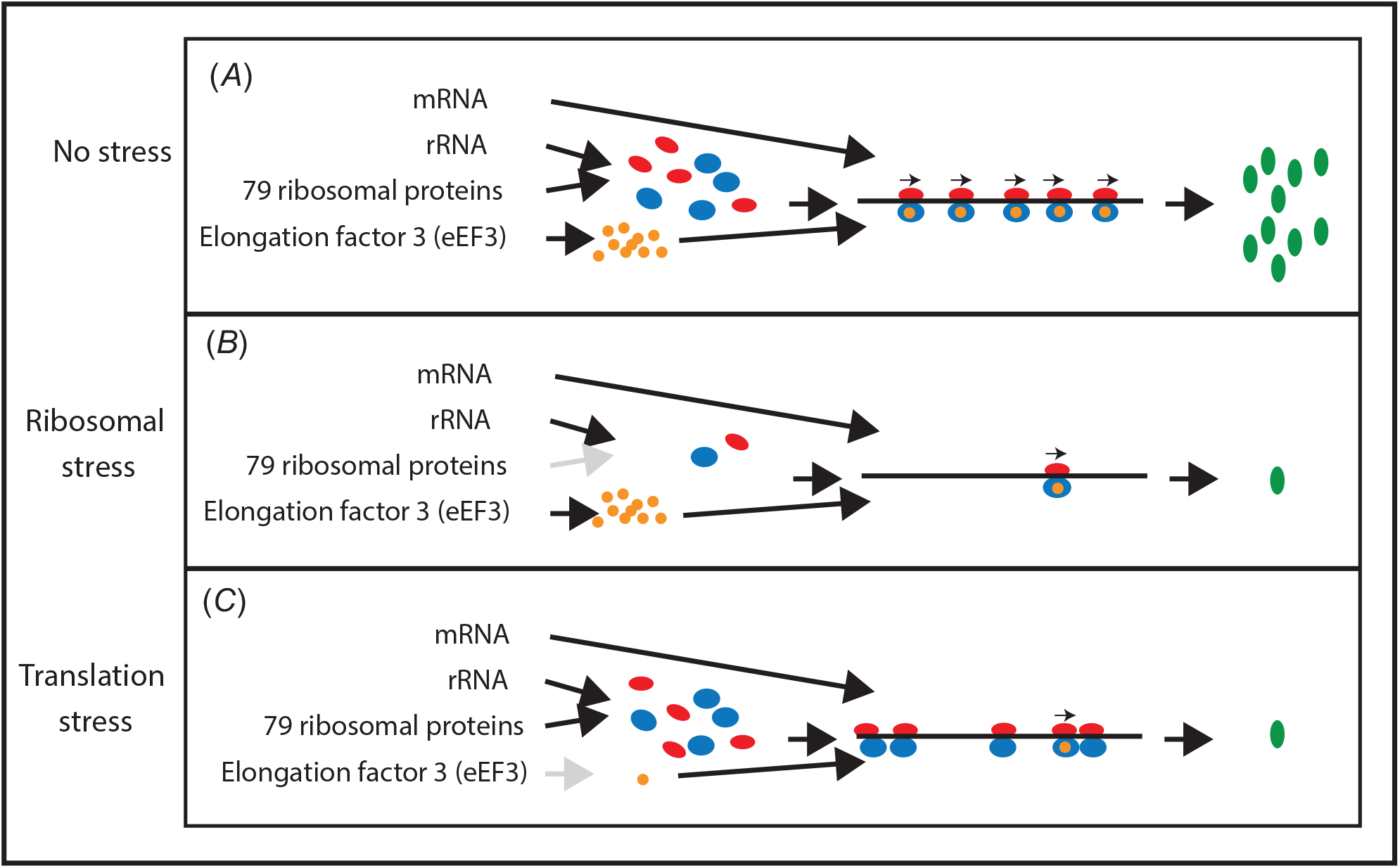
Stress conditions used for RNA-seq analysis. (*A*) Exponential growth with uninhibited protein synthesis. The 40S and 60S ribosomal subunits are shown in red and blue, respectively, eEF3 in orange, and protein products in green. Arrows above the 80S ribosomes on the mRNA indicate elongation-competent ribosomes, each associated with an eEF3 molecule. (*B*) Nucleolar stress: ribosome assembly is inhibited by repression of uL4 synthesis, resulting in reduced numbers of ribosomes relative to cell mass, but the remaining ribosomes are peptide-elongation competent because the number of eEF3 molecules is not decreased. (*C*) Translation stress: the number of ribosomes is unchanged, but ribosomal translocation on the mRNA is inhibited as eEF3 is depleted. See Fig.S1 and (25, 26, 60) for details.

## Results

### Establishing nucleolar and translation stress

To establish nucleolar and translation stress, we used strains in which the only gene for r-protein uL4 or Translation Elongation Factor eEF3 is controlled by the *GAL1/10* promoter inserted at the normal genomic position of the genes *RPL4A* or *YEF3/TEF3* (Table S1). We refer to these strains as Pgal-uL4 and Pgal-eEF3. In galactose medium the *GAL1/10* promoter is active and both strains grow with a doubling time of ∼2.1 hours at 30°C. Sucrose gradient analysis indicated that ribosome assembly and translation are normal: the distribution of free ribosomal subunits, 80S, and polysomes does not differ from wildtype and no abnormal ribosome peaks are seen (Figs. S1A, C, and F) (25, 26).

Shifting Pgal-uL4 from galactose to glucose medium represses the *GAL1/10* promoter and abolishes the synthesis of uL4, blocking ribosome assembly and slowing growth because the assembly of new ribosomes is obstructed (Figs. 2A and S1D). This decrease in the number of ribosomes is independent of whether *RPL4A* (used in all experiments except Fig. S1E) or its paralogue, *RPL4B*, is the source of uL4 protein (compare Figs. S1D and E). In all experiments reported here, except in Fig. S1E, *RPL4B* was inactivated and *RPL4A* was the only source of r-protein uL4. Moreover, abolishing the synthesis of other r-proteins generates the same pattern, suggesting that the consequences of the abolition of uL4 synthesis are representative of nucleolar stress in general (25).

**Fig. 2.**
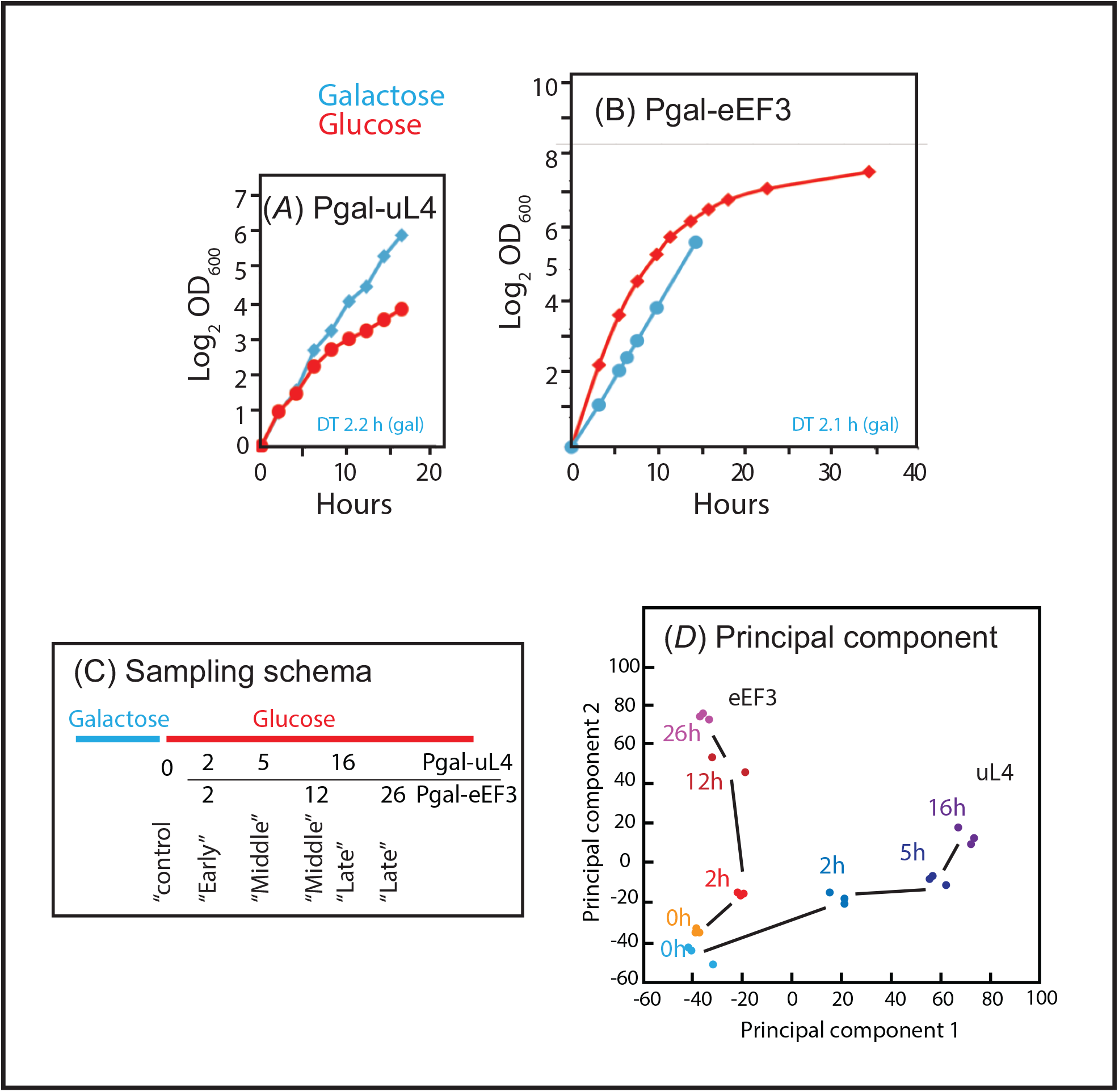
Repression of uL4 and eEF3 synthesis: RNA-seq analysis. (*A-B*) Growth curves of Pgal-uL4 and Pgal-eEF3 growing in galactose (blue) and after a shift from galactose to glucose medium (red). (*C*) Schema for the sampling of cultures. The sampling times were selected to assure that the growth rates (d(OD600)/dt) of the Pgal-uL4 and Pgal-eEF3 cultures are approximately equal at the time of withdrawing the control, early, middle, or late samples. The sampling times of each sample are indicated. (*D*) Principal component analysis of the RNA-seq data in samples taken at the indicated times in the two strains. Results from Pgal-uL4 are shown in blue colors and Pgal-eEF3 results are shown by colors in yellow-to-red colors.

Shifting Pgal-eEF3 to glucose medium decreases the number of eEF3 molecules (26), which slows growth since eEF3 is essential for ribosome translocation but does not affect the cell concentration of ribosomes (Fig. S1F-G). Despite the decreasing synthesis of eEF3, the growth rate initially increases, the result of an upsurge in the differential rate of ribosome formation after the nutritional “shift-up” (Fig. 2B) (6, 27). As the number of eEF3 molecules decreases, the growth rate progressively decreases. No shift-up effect was seen in the growth curve for Pgal-uL4, concurring with the abolition of ribosome production after the shift to glucose medium, which prevents an increase in ribosome number (Fig. 2A). Previous results indicate that the effect of the shift-up on gene expression stabilizes within an hour after the shift (6); we chose 2 hours post-shift-up as the time to begin analyzing the transcriptome in cells subjected to either of the two assaults on the protein synthesis capacity (Fig. 2C).

### General trends of the transcriptome

We collected RNA samples from triplicate-independent cultures of Pgal-uL4 and Pgal-eEF3 before and at the indicated times after shifting from galactose to glucose medium. The sampling times were chosen to assure approximately equal growth rates of the two cultures at the time of sampling, estimated from respective growth curves. We refer to the samples as “control”, “early”, “mid”, and “late” (Fig. 2C). Bio-Analyzer Electrophoretograms showed high rRNA integrity in all but one of the middle samples from Pgal-eEF3, which was therefore discarded. The remaining 23 samples were subjected to RNA-seq analysis; mapping the results to the *S. cerevisiae* genome (SGD; https://www.yeastgenome.org/) yielded 2.3-3.3_*_10^7^ reads from each sample (Table S2A). The abundance of mRNA from specific genes (expression level) in each sample was calculated by mapping reads to each gene/open reading frame in the yeast genome (gene “read counts”) (Table S2B) and the change in the expression of each gene (“fold change”) after the onset of stress was calculated as the read counts in each sample normalized to the read counts in the control sample of the same strain. Table S3A lists the log2 of the fold change (lfc) for 5348 open reading frames all of which had at least 50 read counts at all sampling times.

To ascertain the similarity between biological replicates from a given strain and the variance among depletion samples collected from Pgal-uL4 and Pgal-eEF3, we performed a principal component analysis (PCA) on the read counts per transcript/gene feature after size factor normalization using the DESeq2 package. The replicate samples for a given time point in each strain clustered tightly indicating high reproducibility (Fig. 2D) as is also indicated by the p-values and False Discovery Rates (FDR) for genes going up or down by 2-fold or more (|lfc|≥1) (Table S3A).

After the induction of stress, the results move along different paths in the PCA space (Fig. 2D), indicating that the transcriptome pattern develops differently during the two stress forms. This concords with the distribution of Differentially Expressed Genes (genes with at least two-fold change in mRNA abundance up (up-DEGs) or down (down-DEGs)) and a False Discovery Rate (FDR) <0.05): DEGs represent 12-26% of the total number of genes at different times, but only 21-35% of all DEGs are observed in both strains at a given time. Furthermore, only 6% of genes always have an |lfc| ≥1 (Table 1). Specific DEGs at each time point are listed in Table S4A.

**Table 1.** Number of differentially expressed genes (DEGs) at different times after repressing synthesis of r-protein uL4 (Pgal-uL4) or Translation Elongation Factor eEF3 (Pgal-eEF3). N: number of DEGs in the sample. F: DEGs normalized to the total number of genes.

| Time | Total DEGs |  |  |  |  |  |  |  | Up-DEGs |  |  |  |  |  |  |  | Down-DEGs |  |  |  |  |  |  |  |
| --- | --- | --- | --- | --- | --- | --- | --- | --- | --- | --- | --- | --- | --- | --- | --- | --- | --- | --- | --- | --- | --- | --- | --- | --- |
|  | Pgal-uL4 |  | Both |  | Pgal-eEF3 |  | All |  | Pgal-uL4 |  | Both |  | Pgal-eEF3 |  | All |  | Pgal-uL4 |  | Both |  | Pgal-eEF3 |  | All |  |
|  | N | F | N | F | N | F | N | F | N | F | N | F | N | F | N | F | N | F | N | F | N | F | N | F |
| Early | 334 | 6.2% | 223 | 4.2% | 104 | 1.9% | 661 | 12.4% | 128 | 2.4% | 17 | 0.3% | 22 | 0.4% | 167 | 3.1% | 206 | 3.9% | 206 | 3.9% | 82 | 1.5% | 494 | 9.2% |
| Middle | 535 | 10.0% | 307 | 5.7% | 296 | 5.5% | 1138 | 21.3% | 334 | 6.2% | 41 | 0.8% | 89 | 1.7% | 464 | 8.7% | 201 | 3.8% | 266 | 5.0% | 207 | 3.9% | 674 | 12.6% |
| Late | 703 | 13.1% | 296 | 5.5% | 410 | 7.7% | 1409 | 26.3% | 447 | 8.4% | 76 | 1.4% | 175 | 3.3% | 698 | 13.1% | 256 | 4.8% | 220 | 4.1% | 235 | 4.4% | 711 | 13.3% |
| All | 165 | 3.1% | 116 | 2.2% | 55 | 1.0% | 336 | 6.3% | 4 | 0.1% | 0 | 0.0% | 12 | 0.2% | 16 | 0.3% | 161 | 3.0% | 116 | 2.2% | 43 | 0.8% | 320 | 6.0% |

### Characteristics of the transcriptome after the shift to glucose medium

The ratio of the read counts in the control samples (t=0) was between 0.70 and 1.3 for 79% of the 5348 genes listed in Table S3. Thus, the baseline expression of most genes was similar in the two strains before the onset of stress. Shifting Pgal-uL4 from galactose to glucose medium reduced the pool of uL4A mRNA by 30-to 60-fold and elevated the eEF3 mRNA abundance by <50% (Table S3A). In contrast, shifting Pgal-eEF3 to glucose medium reduced the pool of eEF3 mRNA by 30-to 100-fold, while the abundance of uL4 mRNA is unchanged. Thus, the repression of uL4 and eEF3 mRNA is specific to each of the two strains.

Besides the exposure to either nucleolar or translation stress, mRNA abundance might also be affected by the change of carbon source. To parse the consequences of these different interferences, we first identified genes whose expression changes most differently, or most similarly, using a Rank Value Difference Analysis (28). This involves ranking the lfc values for each gene within each sample, calculating the rank value difference (RVD) for the corresponding genes in early, middle, and late samples from the two strains, and finally ranking the sum of the absolute RVDs for each gene at the three time-points (∑|RVD|; Table S3B). The genes with the highest ∑|RVD| are most likely to show different expression patterns, on average, in the two strains after the induction of stress. Conversely, on average, the genes with the lowest ∑|RVD| are most likely to have similar expression patterns.

#### Many mitochondria-related genes are repressed during both nucleolar and translation stress

Submitting the 200 genes with the lowest ∑|RVD| to the “GO Enrichment Analysis Tool” of the “Gene Ontology” database (http://geneontology.org/) returned several GO terms for genes mostly encoding mitochondrial proteins (Fig. 3). The average ratio between the read counts for these genes in the Pgal-uL4 and Pgal-eEF3 control galactose cultures is 1.2 (Fig. 3, column 1), indicating that their expression levels are similar in the two strains before the imposition of stress. After the switch to glucose medium, most of the genes are repressed in both strains (Fig. 3). Moreover, mRNA from the “Low-affinity glucose transporter of the major facilitator superfamily” *HXT1* and *HTX3* are induced.

**Fig. 3.**
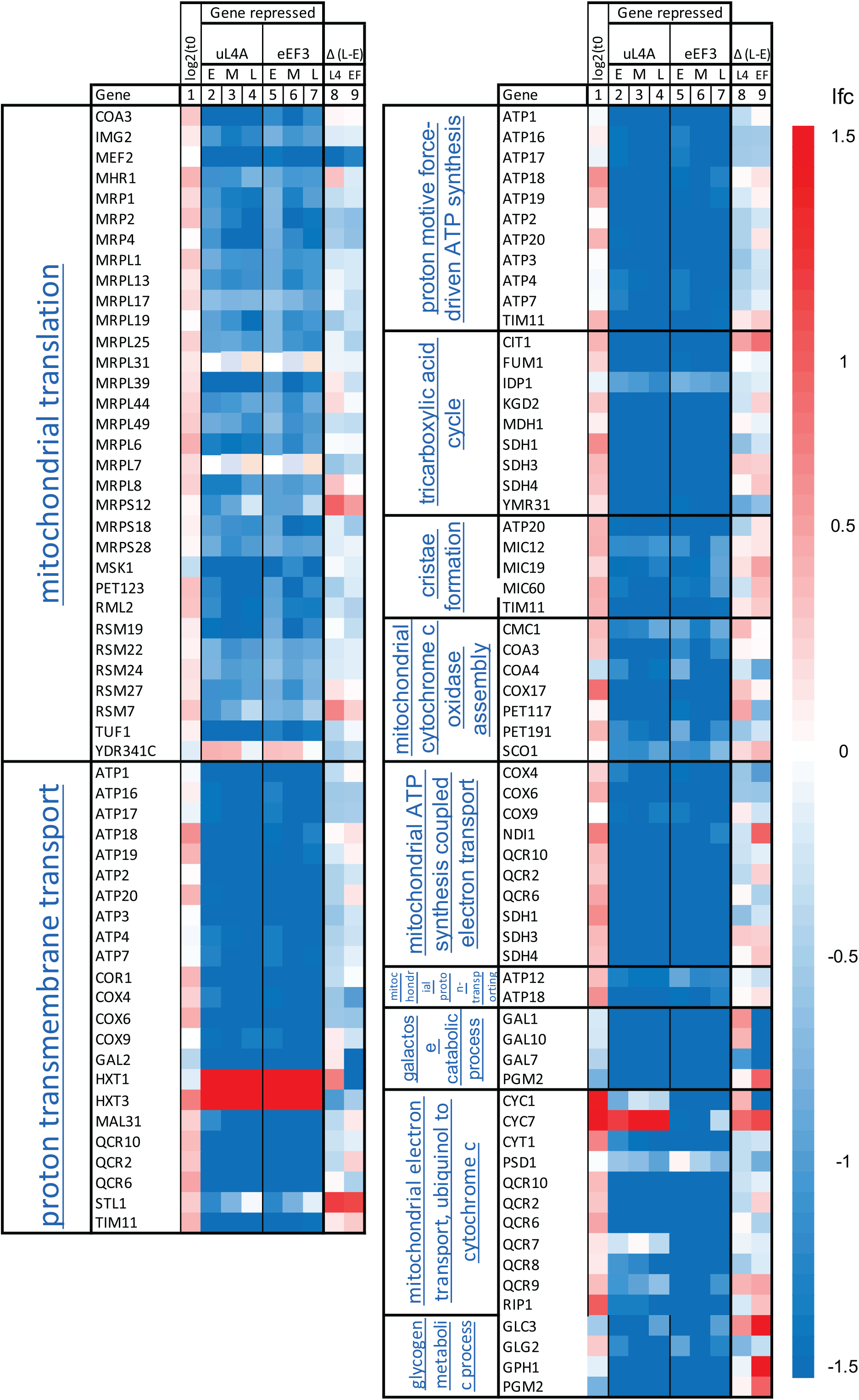
Heatmaps for genes with the most similar change of expression (lfc) after the shift from galactose to glucose medium. Submission of the 200 genes with the lowest Rank Value Difference (Table S3B; see also text) to GO enrichment analysis at http://geneontology.org/ returned the Biological Process GO-terms (False Discovery Rates (FDR) in parenthesis): mitochondrial translation (4E-12), proton transmembrane transport (1E-3), proton force-driven ATP synthesis (4E-3), tricarboxylic cycle (4E-2), cristae formation (4E-3), mitochondrial cytochrome c oxidase assembly (1E-2), mitochondrial proton-transporting ATP synthesis complex assembly 2E-2), galactose metabolic process (2E-2), mitochondrial metabolic process, ubiquinol to cytochrome c (4E-2), and glycogen metabolic process (5E-2). The heatmaps show lfc for genes listed collectively under these GO terms. The first two columns show GO terms (http://geneontology.org/) and the gene names (https://www.yeastgenome.org/), respectively. Column 1 shows log_2_ (read count in Pgal-uL4/read count in Pgal-eEF3) at t=0. Columns 2-4 show the lfc (log_2_ of fold change) in Pgal-uL4 gene expression (read count for the early (E), middle (M), and late (L) samples, respectively, relative to Pgal-uL4 read count at t=0 control). Columns 5-7 similarly show the change in Pgal-eEF3 gene expression in the E, M, and L samples relative to the control culture. Columns 8 and 9 show the difference between lfc in the L and E samples for Pgal-uL4 and Pgal-eEF3, respectively, i.e., the change in gene expression between the late and early time points in each strain. The scale for the heatmaps is shown to the right. The heatmap scale from -1.5 to +1.5 is shown to the right.

We conclude that the pattern of expression of genes with the lowest ∑|RVD| suggests that they are under glucose repression, in accord with the previous analysis of the regulation of genes for the formation of mitochondria (29). Note that the lfc for the mitochondrial genes changes little after 2 hours, indicating that the glucose repression is established and stable within the first 2 hours after the media shift.

#### The abundance of mRNA for r-proteins changes differently during nucleolar and translation stress

Submitting the 200 genes with the highest ∑|RVD| to the GO term enrichment yielded a single GO term, cytoplasmic translation, which is enriched by 3.4-fold (FDR 2.2E-02). This prompted us to generate heatmaps of all genes encoding r-proteins, ribosome assembly (Ribi) factors, and ribosome translation factors (Figs 4-5). The mRNAs for these genes account for about 20%, 3%, and 3% of all mRNAs, respectively, in the galactose control cultures (t=0) for both strains. Furthermore, the average ratio of the read counts Pgal-uL4 and Pgal-eEF3 before the imposition of stress is between 0.8 and 1.2, indicating similar baseline expression in the control cultures (columns 1 in Figs. 3-6 and Fig. S3).

**Fig. 4.**
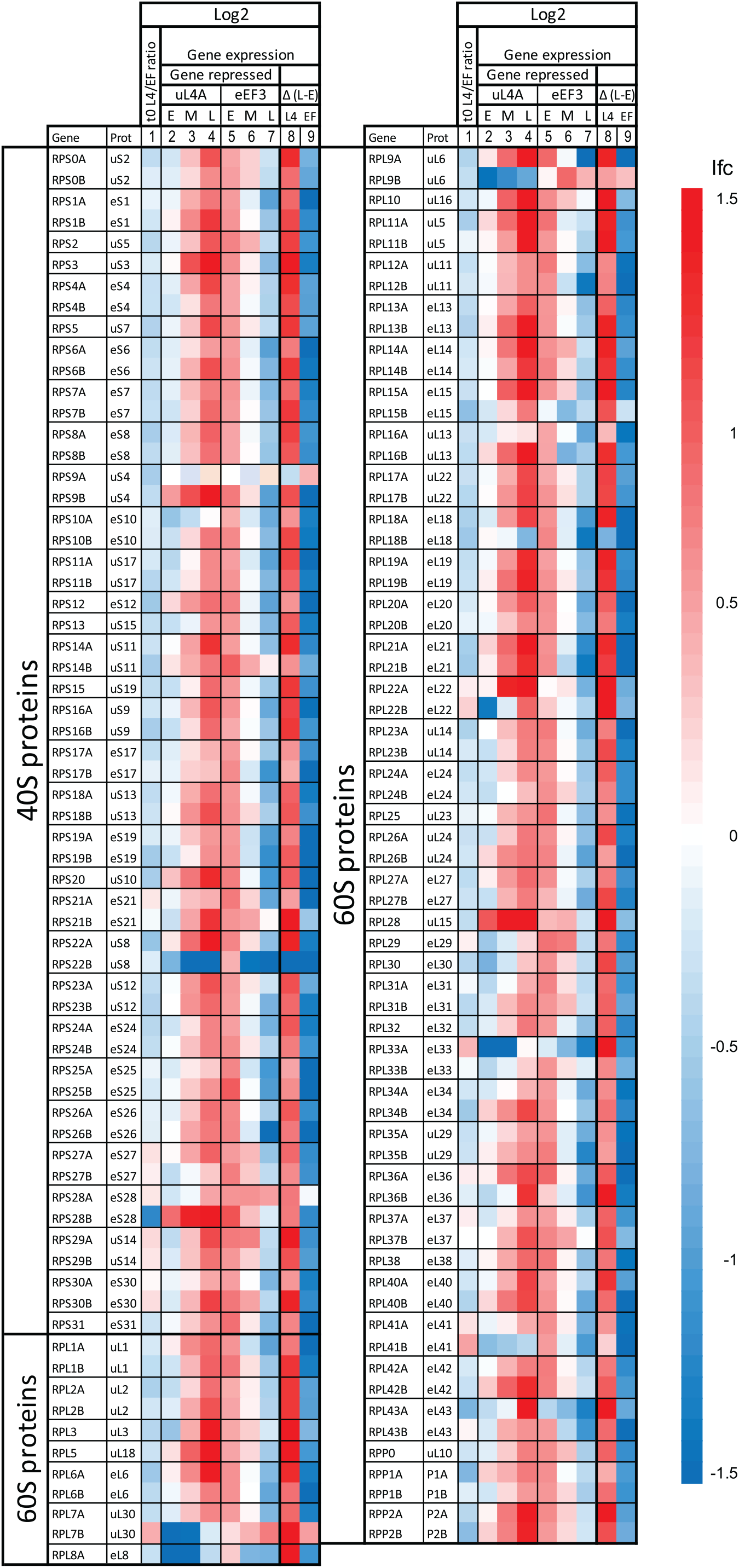
Heat maps for genes for the 60S and 40S cytoplasmic ribosomal proteins during nucleolar and translation stress. The two left columns show gene names (https://www.yeastgenome.org/) and the names of their encoded proteins according to the nomenclature of Ban et al (61). The heatmaps are arranged as explained in the legend in Fig. 3.

**Fig. 5.**
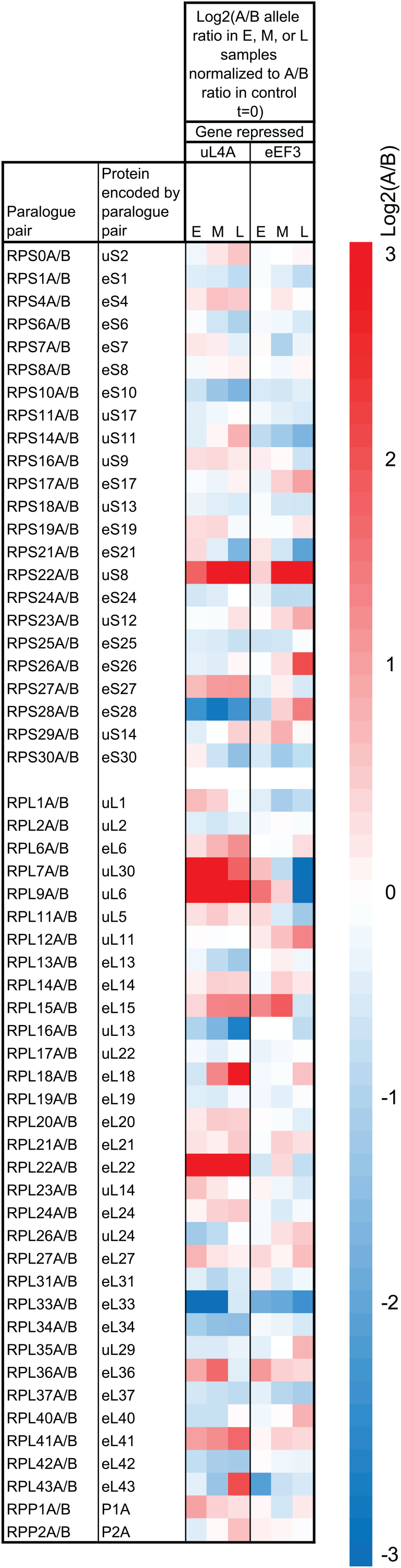
Heat maps for the relative abundance of mRNA from the A and B paralogous alleles encoding the same r-proteins. The heat map shows the ratio between the A and B alleles for a given r-protein normalized to the corresponding ratio in the control sample. Note that the scale of the heat maps goes from -3 to +3.

Ribosomal precursor particles are transported through the nuclear pores on their way to the cytoplasm where the final maturation into translation-competent ribosomes occurs. Hence, we analyzed the expression of mRNA for nuclear pore proteins. The mRNAs for these proteins are also not co-regulated with the r-proteins transcriptome during nucleolar and translation stress (Fig. 6). We conclude that none of the genes that support ribosome biogenesis and function are coordinated with the r-protein mRNA abundance during interference with the functions they support.

**Fig. 6.**
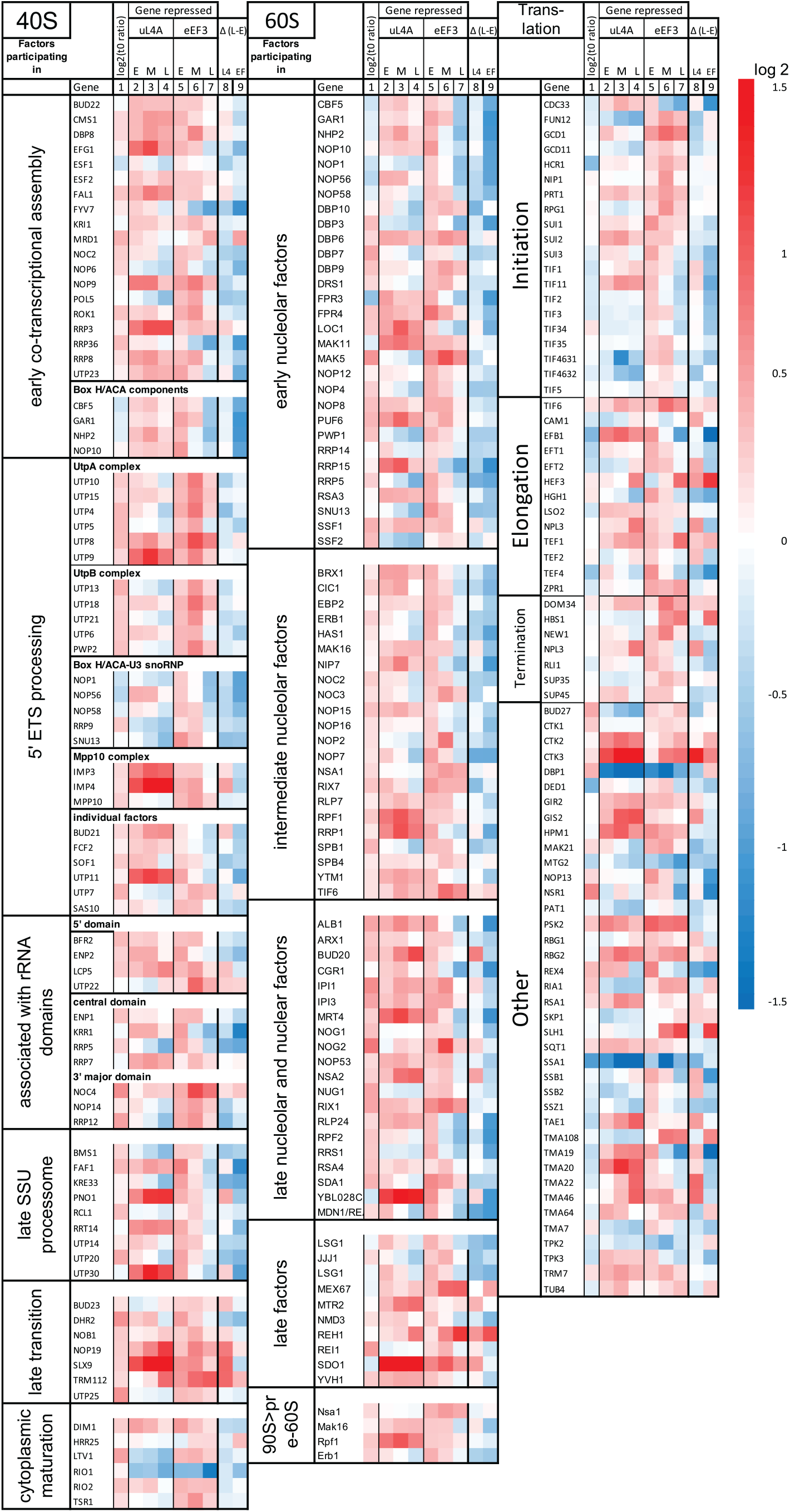
Heatmaps for the 60S and 40S assembly factors and translation factors during nucleolar and translation stress. The columns are organized as explained in the legend to Fig. 3. The genes are ordered according to their function in the assembly of each subunit or ribosome translation function (4, 62, 63).

**Fig. 7.**
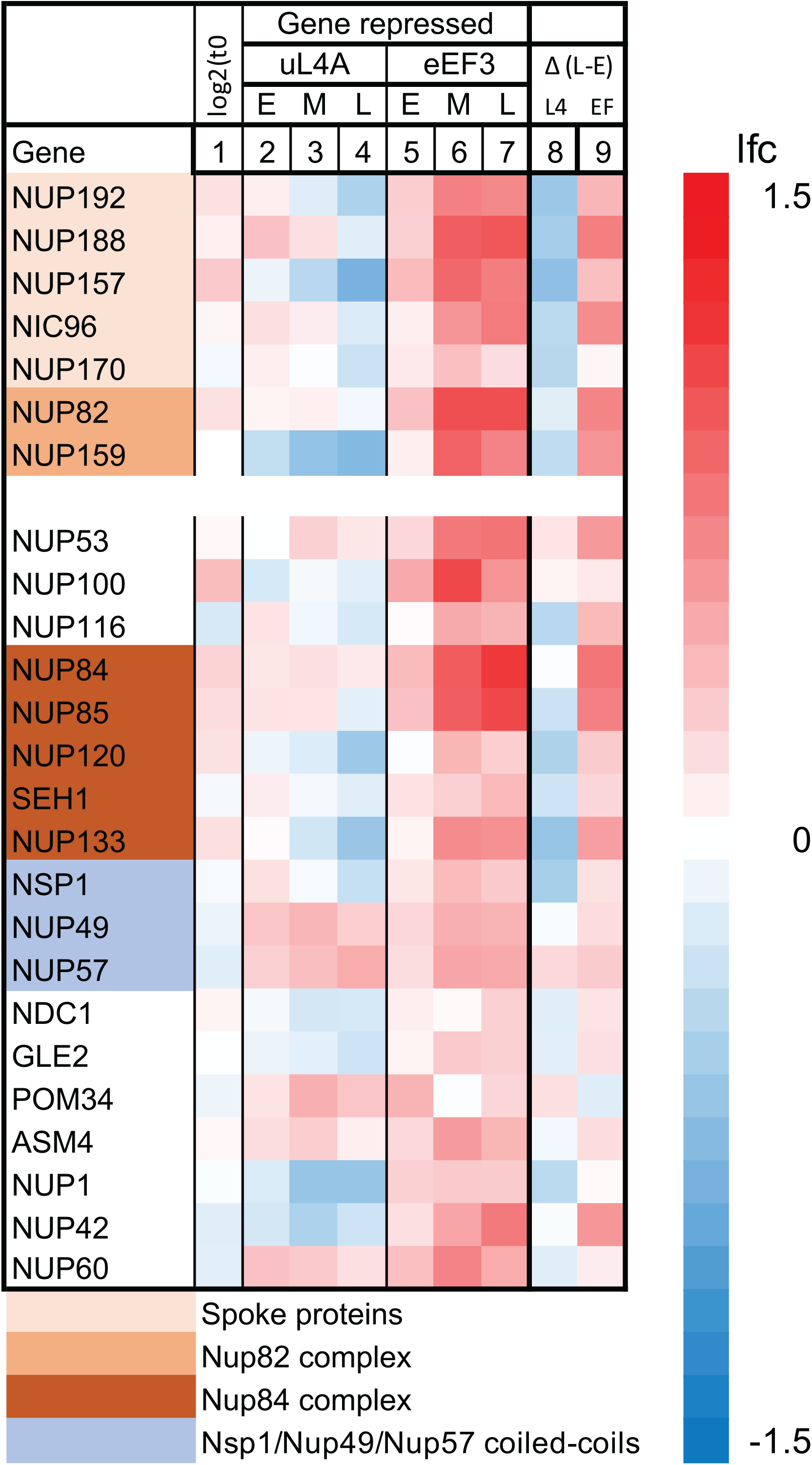
Heatmaps for genes encoding nuclear pore proteins. The columns are organized as described in the legend to Fig. 3. Genes encoding components of subcomplexes of the nuclear pore are color-coded (64).

Interestingly, r-protein mRNA abundance is regulated in opposite directions during nucleolar and translation stress. During the repression of uL4 synthesis, mRNAs for other r-proteins are initially repressed, then switch to induction after 2 hours (Fig. 4, columns 2-4). Repression of eEF3 synthesis results in the opposite pattern: an initial induction as expected from the nutritional shift-up followed by repression (Fig. 4, columns 5-7). Importantly, as opposed to the genes for mitochondrial proteins, the lfc for r-protein genes keeps changing after 2 hours, indicating that the r-protein genes are regulated by the imposed stress rather than the media shift (see columns 8 and 9 in Fig. 4).

The lfc values for some samples have FDR >0.05 since they are close to 0. Consequently, we plotted all lfc values against time, which shows that essentially all lfc values fall on curves with positive slopes after inhibiting uL4 synthesis and negative slopes after repressing eEF3 synthesis (Fig. S2). The only exception is the values for RPS22B, which decline after abrogating uL4 synthesis. Overall, we conclude that the mRNA for r-protein genes is induced during nucleolar stress but repressed during translation stress.

#### The relative expression of the paralogues for some r-proteins changes during stress

About two-thirds of the *S. cerevisiae* r-proteins are encoded by pairs of paralogous genes, called A and B, whose protein products differ by a few amino acids. To determine if the ratio of paralogous messengers changes during stress we calculated the ratio of read count for the A and B alleles normalized to the corresponding ratio in the control cultures. Note that we cannot determine the absolute ratio between the expression of the paralogous allele as we do not know if the RNA-seq procedure registers both alleles with the same efficiency, but this uncertainty is eliminated by normalizing the data to the control sample (t=0). The results show that the relative expression of paralogue pairs changes more than 2.5-fold for uS8, eS28, uL6, eL18, eL22, uL30, and eL33, but less than 50% for most other r-protein paralogue pairs (Fig. 5).

#### Genes for ribosome assembly and ribosome translation do not change coordinately with r-protein genes or with each other

Ribosome homeostasis and function depend on hundreds of auxiliary factors that support ribosome formation and mRNA translation. Unexpectedly, the mRNA abundance for neither the ribosome assembly factors (Ribi) nor the translation factors is coordinated with the changes in the r-protein mRNAs (Fig. 6). The lfc for most of these mRNAs change less than 2-fold and show no obvious common pattern. The mRNAs for nuclear pore proteins are also not coordinated with r-protein mRNA, even though about 1000 precursor ribosomes are exported through the nuclear pores from the nucleus to the cytoplasm each minute during exponential growth (30).

#### Other GO terms and gene groups

Genes with consecutive ∑|RVD| ranks between the top- and the bottom-ranked genes (Table S3B) values fail to return statistically significant GO terms. A search for other patterns among DEGs showed that GO terms defined by DEGs at specific time points in each strain mostly include genes for metabolic functions while others are closely related to GO terms for glucose-repressed genes mentioned above (Table S4).

We also searched for genes with different combinations of increasing and decreasing lfc trends during stress. GO term Enrichment Analysis of the resulting lists of genes generated GO terms for metabolism genes, protein transport, and mitotic cell cycle (Table S5), suggesting that significant parts of the cell physiology change during nucleolar and translation stress.

## Discussion

Interference with the translation machinery’s formation or function often has serious consequences for an organism’s fitness, resulting in various human diseases. It is assumed that insufficient translation capacity is a significant contributor to the disease state, but it is not clear whether reduced translation capacity as such, no matter the cause, always has the same effect on the function of the cell. To answer this question, we compared the yeast transcriptome after abolishing ribosome biogenesis (nucleolar stress) and blocking ribosome translocation (translation stress). Both stresses inhibit protein synthesis, cell growth, and cell cycle progression, but it is unknown if the transcriptomes associated with these calamities are the same or different. Our results show that both the r-protein mRNA abundance and paralogue composition of mRNAs are regulated differently during the two stresses.

### Regulation of ribosomal protein mRNA abundance during stress

Both the abundance and composition of r-protein mRNA are regulated differently by the two forms of stress. The abrogation of ribosome assembly by repressing the synthesis of uL4 prevents the assembly of the other newly synthesized proteins into stable, functional ribosomal particles (25). The disjunction of ribosome assembly initially decreases r-protein mRNA abundance (Fig. 4), presumably due to the accumulation of insoluble cytotoxic precipitates containing r-proteins and other proteins that directly or indirectly inhibit the transcription of r-protein genes (31). Interestingly, two hours after the blocking of ribosome assembly, r-protein mRNA abundance increases (Fig. 4 and Fig. S2). We can think of two explanations: Since it is known that free r-proteins are attacked by proteasomes, proteasome activity may increase, thereby reducing the pool of free r-proteins (32, 33). Second, the chaperone-mediated transport of nascent r-proteins to the nucleus may increase, which would stabilize the r-protein mRNA (34).

The initial increase in r-protein mRNAs after stopping eEF3 synthesis is likely due to the nutritional shift-up caused by transferring the culture to glucose medium. Nevertheless, two hours after inducing translation stress, the r-protein mRNA abundance is repressed through the rest of the experiment. Perhaps as the number of eEF3 molecules falls below the number of ribosomes, the wait time for an eEF3 factor increases, temporarily stopping the movement of a fraction of the ribosomes and resulting in collisions between mobile and stationary ribosomes (Fig. 1A(iii)). This, in turn, may trigger mRNA degradation (35, 36).

The difference in the r-protein transcriptome during the two types of stress correlates with the ratio between cell mass and ribosome number. During nucleolar stress, the ribosome formation is stopped, but the pre-existing ribosomes keep synthesizing proteins, i.e., the cell-mass-to-ribosome ratio increases. During translation stress, the ratio stays constant because the synthesis of all proteins, including r-proteins, declines at the same rate as ribosome translocation is inhibited. Thus, the r-protein mRNA abundance is upregulated only when the ratio between the growth rate and the ribosome number increases. It may also be important to note that the nucleolar stress is directed toward a nucle(o)lar process, while the translation stress is directed towards a cytoplasmic process.

### Differential expression of mRNA for r-protein paralogues

More than two-thirds of the *S. cerevisiae* r-proteins are encoded by duplicated genes (A and B paralogous alleles) whose gene products can differ by a few amino acids. It is well established that the composition of paralogous r-proteins in individual ribosomes often varies in response to developmental or other external stimuli, thereby potentially generating “specialized ribosomes” with altered translational properties. However, a consensus on the biological significance of ribosome heterogeneity has not yet emerged (37-48).

To determine the effects of nucleolar and translation stress on the accumulation of mRNAs for paralogous r-proteins, we calculated the ratio between the mRNA from the A and B paralogues changes during stress, normalized to the ratio in the control cultures. Only minor changes in the paralogue ratio occurred for most r-proteins (Fig. 4). However, the relative amount of mRNA from paralogues encoding eS28, uL6, eL18, eL22, uL30, and eL33 changed by 2.5-fold or more during one or the other form of stress, while uS8 responded similarly to both stresses (Fig. 4). Since the ratio changed differently during the two types of stress for all proteins mentioned, except uS8, the change in paralogue compositions is due to the imposition of stress, **not** the change in the growth medium. This is compatible with the notion that the r-protein composition of individual ribosomes might mechanistically contribute to adapting the cell physiology during changing growth conditions.

### Genes encoding factors that support ribosome biogenesis and function are not co-regulated with r-protein genes

The ribosome assembly factors (Ribi) and translation factors are essential for the formation and function of ribosomes, respectively. Previous investigations have suggested that the Ribi and r-protein genes are both parts of a large network regulated by the Sfp1 transcription factor (49, 50). However, our results show that the genes for neither the Ribi nor the translation factors are coregulated with the r-protein mRNA during either ribosome or translation stress (Fig. 5), demonstrating that the coregulation of Ribi and ribosome auxiliary factors with r-proteins is not obligatory. We also note that the expression of Ribi and r-protein genes are not coordinated during the inhibition of rRNA synthesis (31).

### Cell cycle arrest

We previously showed that blocking either ribosome biogenesis or peptide chain elongation results in the accumulation of cells that have completed cytokinesis, but not cell separation (26). To investigate the gene expression associated with this phenomenon, we searched for mRNAs encoding proteins that contribute to septum breakdown and found that mRNA for several genes under the GO term “septum digestion after cytokinesis” are repressed during both nucleolar and translation stress. For example, Egt2 mRNA, encoding a cell wall glucanase, and Dse4 mRNA, encoding a daughter cell-specific secreted protein similar to glucanases, are downregulated (Table S6A) (https://www.yeastgenome.org/). We surmise that the downward trend of mRNAs for septum digestion may contribute to cell cycle arrest during both nucleolar and translation stress.

To determine if other GO term genes are repressed in response to either type of stress, we performed a GO term enrichment analysis of other genes with a Δlfc ≤1. This identified the GO terms of “mitotic nuclear division”, “cell division”, and positive cell cycle regulation during the repression of uL4 synthesis, but repressing eEF3 synthesis did not enrich for the same GO terms, indicating that the expression of these genes is not coordinated with the inhibition of the cell cycle progression (Table S5). However, examination of mRNA abundance for genes whose expression spikes during specific phases of the mitotic cell cycle during unrestrained growth (http://www.cyclebase.org/) showed that mRNA from histone genes is repressed during both nucleolar and translation stress (Table S6B), which concords with the finding that ribosome biogenesis is required for passage through Start (G1 to S transition) (51, 52).

### Other mRNAs of physiological importance

A search for genes with an |Δlfc|≥1 after the onset of nucleolar or translation stress identified several additional GO terms that relate to other physiological functions not already discussed. These include “Cellular response to oxidative stress”, “Regulation of chromosomal organization”, “Response to abiotic stimulus”, “Response to extracellular stimulus”, and “Protein transport to vacuole involved in ubiquitin-dependent protein catabolic process via the multivesicular body sorting pathway” (Table S5). These findings support the idea that ribosome formation and function are central to the regulation of cell growth and resource use.

### Implications

Our data show that the transcriptome evolves differently after inhibiting ribosome formation (nucleolar stress) and impeding ribosome translocation (translation stress), showing that the stress response to a reduction of the protein synthesis capacity is not determined simply by the reduction of protein synthesis, but also by the **process** that lowers the translation capacity. We suggest that a given mutation simultaneously affects **both** the synthesis of ribosomes **and** the ribosome function. This notion is supported by the different human phenotypes resulting from the coding region and 5’UTR mutations in the genes for r-protein uL6 (*RPL9*) (47). In conclusion, our transcriptome analysis suggests that a given mutation can generate a unique combination of ribosomal biogenesis and translation responses that in turn generates a unique transcriptome and cell fate for each mutation.

## Materials and Methods

### Strains and growth conditions

Strains are listed in Table S1. Where indicated, the gene for r-protein uL4 or Translation Elongation Factor 3 (eEF3) is under the control of the *GAL1/10* promoter. We refer to these strains as Pgal-uL4 and Pgal-eEF3, respectively.

Cultures were grown asynchronously in YEPGal media (1% yeast extract, 2% peptone, and 2% galactose) at 30°C until mid-log phase (OD_600_=0.8–1.0 corresponding to 1.5–2 × 10^7^ cells/ml) and were then shifted to YPD medium (1% yeast extract, 2% peptone, and 2% glucose). All experiments were done in biological triplicates. Cultures were diluted as necessary with pre-warmed media to keep the OD_600_ <1.0. We did not expose the cultures to cycloheximide before harvest because this treatment can influence both polysome content and mRNA abundance (53).

### RNA preparation and analysis

Samples (1 OD unit) were taken from cultures before and at the indicated times after the shift to glucose medium (Fig. 2C). Total RNA was extracted using the Ribopure Yeast kit (ThermoFisher, USA) following the manufacturer’s protocol. Ribosomal RNA integrity was checked by Bio-Analyzer (Agilent Technologies, USA). One of the three Pgal-eEF3 12h samples did not pass quality control and was not sequenced. RNA samples were enriched for mRNA using poly-A attached magnetic beads followed by Paired-end RNA-seq library preparation using the TruSeq RNA Sample Prep kit (Illumina). One hundred nucleotides were sequenced using the HiSeq platform (Illumina) from both ends of each cDNA; sequence reads were aligned against the *Saccharomyces cerevisiae* S288c reference genome using the Tophat2 aligner (54). Sequence annotation and read counts per gene were generated using HTseq (55) and Subread (56) and compared for consistency. We obtained a mean of 2.3-3.1_*_10^7^ and 2.5-3.3_*_10^7^ reads mapping to the *S. cerevisiae* BY4741 genome for each sample of Pgal-uL4 and Pgal-eEF3, respectively (Supplemental Table S2). DESeq2 (57) was used for calculating differential gene expression between RNA samples relative to the 0-hour sample. A gene was classified as a differentially expressed gene (DEG) if the false discovery rate (FDR) value was <0.05 and the absolute log_2_ (fold change) value was ≥1. The Python Sci-kit-learn was used for Principal Component Analysis. Heatmaps were generated by Excel software. The LFC and FDR of all genes with <50 reads at all times are in Tables S3A.

**Gene ontology enrichment analysis** was performed using the Gene Ontology Resource (http://geneontology.org/); Release Date 7-1-22

### Identification of genes expressed most and least differently during nucleolar and translation stress

To identify the genes that were regulated most and least differently during nucleolar and translational stress we ranked genes by fold change of expression for each time point and calculated the Rank Value Difference (RVD) by subtracting the gene rank values for individual genes in the Pgal-eEF3 samples from the rank values of the corresponding Pgal-uL4 samples. The absolute and actual RVD values for each gene were summed over three time points to calculate the absolute rank-sum and rank-sum, respectively (Table S3B). Finally, the actual rank-sum was subtracted from the absolute value of the rank-sum to calculate the rank sign of each gene (Table S3C). Genes with the highest absolute rank-sum values differ the most in the fold change between the two strains (Pgal-uL4 and Pgal-eEF3) over the three-time points of treatment. The genes with high-rank sign values are the genes that had a higher rank during repression of uL4 synthesis relative to repression of eEF3 synthesis at a specific time point but changed to a lower rank relative to eEF3 repression at other time points and vice versa.

## Data availability

The raw sequencing reads from this study have been submitted to the NCBI sequence read archive (SRA) under BioProject accession no. PRJNA693823. The data are also available upon request.

## Acknowledgments

This work was supported by grant 0920578 from the National Science Foundation to JMZ and LL. Additional funding was provided by an internal appropriation from the University of Maryland, Baltimore County to LL (no grant number). We thank B. Traasdahl for help with the manuscript.

## Legends to Supplementary Tables

**Table S1**. Strains used

**Table S2**. Analysis of reads. (A) The number of total reads in individual samples. (B) Read counts for individual genes for which the average number of reads is ≥50 at all times (5348 genes). The last column lists the read count at t=0 in Pgal-uL4 divided by the corresponding number in Pgal-eEF3.

**Table S3**. Analysis of mRNA abundance per gene, strain, and time. (A) lfc, p, and FDR (False Discovery Rate) values are listed for all genes for which all samples had ≥50 reads (5348 genes). Feature.ID and Gene name used are according to the Saccharomyces Genome Database (https://www.yeastgenome.org/). Where no gene name has been assigned, the Feature.ID was used as the gene name. The RPL4A and YEF3/TEF3 are shown in red to indicate that these genes are repressed in the Pgal-uL4 and Pgal-eEF3 strains, respectively. Note that p values for some genes are below the lowest value that can be shown in an Excel cell. The original results for these samples were listed as 0.0E+00. We have arbitrarily changed this to 1E-300. Colors serve as visual help to distinguish columns only. (B) lfc values are sorted according to the absolute sum of Rank Value Differences (RVD); see text. (C) lfc values sorted according to Rank Sign; see text.

**Table S4**. Differentially expressed genes (DEGs) at different times after abrogating uL4 or eEF3 synthesis. (A) Genes with lfc≥1 and FDR≤0.05 are shown for each sample. (B) Biological process GO terms found using DEGs from early samples as input. (C) Biological process GO terms found using DEGs from middle samples as input. (D) Biological process GO terms found using DEGs from late samples as input. The heat maps are arranged as in Fig. 3. GO term searches were done using the GO Enrichment Analysis in the Gene Ontology Resource (http://geneontology.org/)

**Table S5**. Analysis of gene expression according to lfc trends after abrogating uL4 or eEF3 synthesis. (A) GO terms identified among genes with |lfc| changing by ≥1 after repressing uL4 and/or eEF3 synthesis. (B) Heatmaps of DEGs under the GO terms identified in (A) using genes with lfc≤-1 as input. The ribosomal protein mRNAs are removed from the GO term Nucleolus but are shown in Fig.4. (C) Heatmaps of DEGs under the GO terms identified in (A) using genes with lfc≥1 as input. The heat maps are arranged as in Fig. 3. GO term searches were done using the GO Enrichment Analysis in the Gene Ontology Resource (http://geneontology.org/).

**Table S6**. Analysis of mRNA abundance from cell-cycle-related genes. (A) Heat maps for genes under the GO term Septum Digestion after Cytokinesis. (B) lfc values for genes with peak expression in specific cell cycle phases. The heat maps are arranged as in Fig. 3.

## Legends to Supplemental Figures

**Fig. S1**. Analysis of ribosomes, uL4 mRNA, and eEF3. (A-G) Sucrose gradient analysis of whole cell extracts. (A-B) Parent strain (BY4741) growing in galactose and glucose. (C-D) Pgal-uL4 (*RPL4A*) growing in galactose and harvested before and 7 hours after shifting the culture to glucose. (E) Pgal-uL4B (*RPL4B*) was harvested 15 hours after shifting from galactose to glucose. The asterisks indicate peaks of “half-mers” (an mRNA with N ribosomes plus an initiating 40S waiting for a 60S subunit), which are enlarged due to the addition of cycloheximide a few seconds before harvest (53). All other cultures were harvested without cycloheximide. (F-G) Pgal-eEF3 growing in galactose and harvested before and 16 hours after shifting the culture to glucose. (H) Northern blot analysis of uL4 mRNA in the parent (BY4741), carrying the empty Pgal cloning vector (pGilda, Clontech), or Pgal-uL4 before and at the indicated times after shifting the culture from galactose to glucose. The difference in length of the mRNA in the two strains is due to substituting the *RPL4A* 5’ UTR in the parent with the *GAL* 5’UTR in the Pgal-uL4 strain. The northern blot was probed with ^32^P-labeled cDNA made by first PCR amplifying the coding region of RPL4B and using the product as template for extension of the 3’ primer in the presence of α-32P dATP. Note that this probe hybridizes to both uL4A and uL4B mRNA due to the high similarity of the coding sequence in the two genes. The *RPL4B* was disrupted and therefore not expressed. (I) Western analysis of eEF3 in Pgal-eEF3 harvested before and at the indicated times (hrs) after shifting the culture from galactose to glucose medium. The blot was probed with Polyclonal anti-eEF3 (1:10,000 dilution) purchased from Kerafast Inc, Boston, MA, USA. Strains with r-proteins eL43 and uS4 under control of the *GAL1/10* promoter (Pgal-eL43 and Pgal-uS4) were used as a control to verify that obliterating ribosome assembly does not affect the synthesis of eEF3 protein (encoded by TEF3 alias YEF3). (J) Growth curve for Pgal-uL4B. Some panels were taken or modified from refs (25) (Panels A-B) and (26) (Panels F, G, and I).

**Fig. S2**. Analysis of the influence of FDR on the kinetics of lfc for individual r-protein genes after abrogating uL4 and eEF3 synthesis. (A) Log_2_(fold change) for individual genes in early, middle, and late samples. Genes for which ≤1 sample has an FDR>0.05 in the two strains, collectively, has an FDR≤0.05 are shown with colored circles and triangles. Genes for which >1 sample has an FDR>0.05 are shown as lines. The data for *RPS22B* are identified (see text). **(B)** Scatter plot comparing the mean lfc for genes having ≤ 1 sample with FDR≤0.05 and genes for which >1 sample is >0.05.

**Fig. S3**. Comparison of mRNA abundance for different classes of genes in the Pgal-uL4 and Pgal-eEF3 control cultures. Box and whisker plots of the ratio between read counts found before the media shift in the two strains. The boxes show the two middle quartiles, and the whiskers show the top and bottom quartiles. The points show statistical outliers.

